# The dynamics of oligodendrocyte populations following permanent ischemia promotes long-term spontaneous remyelination of damaged area

**DOI:** 10.1101/2023.12.18.572134

**Authors:** Gerardo Martín-Lopez, Paula R. Mallavibarrena, Mario Villa-Gonzalez, Noemi Vidal, Maria José Pérez-Alvarez

## Abstract

Stroke is a major public health concern, whit limited clinically approved interventions available to enhance sensorimotor recovery beyond reperfusion. Remarkably, spontaneous recovery is observed in certain stroke patients, suggesting the existence of a self-brain repair mechanism not yet fully understood. In a rat model of permanent cerebral ischemia, we described an increase in oligodendrocytes expressing 3RTau in damaged area. Considering that restoration of myelin integrity ameliorates symptoms in many neurodegenerative diseases, here we hypothesize that this cellular response could trigger remyelination. Our results revealed after ischemia an early recruitment of OPCs to damaged area, followed by their differentiation into 3RTau^+^ pre-myelinating cells and subsequent into remyelinating oligodendrocytes. Using rat brain slices and mouse primary culture we confirmed the presence of 3RTau in pre-myelinating oligodendrocytes and a subset of mature. The myelin status analysis confirmed long-term remyelination in the damaged area. Postmortem samples from stroke subjects showed a reduction in oligodendrocytes, 3RTau^+^ cells, and myelin complexity in subcortical white matter. In conclusion, the dynamics of oligodendrocytes populations after ischemia reveals a spontaneous brain self-repair mechanism which restores the functionality of neuronal circuits long-term by remyelination of damage area. This is evidenced by the improvement of sensorimotor functions in ischemic rats. A deep understanding of this mechanism could be valuable in the search for alternative oligodendrocyte-based, therapeutic interventions to reduce the effects of stroke.

## Introduction

Stroke is one of the most common debilitating diseases. Characterized by high mortality, it ranks as the second leading cause of death worldwide and it is a major contributor to long-term disability (Campbell *et al*., 2019). Reperfusion is currently the only approved therapeutic intervention available for this condition. However, this procedure is not suitable for all patients, and it increases the risk of hemorrhagic transformation, thus promoting cerebral ischemia–reperfusion injury (Zhang *et al*., 2023). Some stroke patients experience spontaneous neurological improvements after damage. This observation supports the notion that brain remodeling can occur after stroke (Jia *et al*., 2019). In this context, unraveling the molecular and cellular mechanisms triggered by the brain to maintain certain functions will contribute to identifying new therapeutic strategies to improve symptoms in ischemic patients.

Oligodendrocytes (OLGs) are particularly vulnerable to ischemia (Hernández *et al*., 2021). These cells can be categorized into distinct populations on the basis of their level of differentiation, morphology, and function in the adult brain (Marques *et al*., 2016). Oligodendrocyte precursor cells (OPCs), identified by nerve/glial-antigen 2 (NG2) expression, are widely distributed throughout the adult CNS and account for the largest population of dividing cells in the healthy brain. They play a crucial role in the lifelong renewal of mature OLGs. OPCs first undergo differentiation into pre-myelinating OLGs before fully acquiring the mature myelinating phenotype (Hernández *et al*., 2021). Mature OLGs provide the rapid saltatory conduction of action potential and ensure protection and nutritional support to axons (Dewar *et al*., 2003). Mature OLGs in the adult brain comprise a heterogeneous population, exhibiting morphological and functional differences that are not only dependent on the brain region but also vary throughout the course of diseases (Frazier *et al*., 2023; Marques *et al*., 2016). The characterization of these OLG subtypes is essential for understanding their role in normal CNS myelination and repair processes. Additionally, such knowledge may offer insights into potential therapeutic strategies for demyelinating diseases, such as stroke, where the dysfunction of OLGs results in impaired neuronal function and contributes to neurodegeneration (Dewar *et al*., 2003; Shi *et al*., 2015).

The remyelination of injured axons is a complex key repair process for restoring functional circuits and maintaining brain function; however, it is not yet fully understood (Franklin and Ffrench-Constant, 2008). White matter alterations have been described in many neurodegenerative diseases, mental disorders, and cerebral ischemia (Aboul-Enein *et al*., 2003). The restoration of myelin integrity ameliorates behavioral, motor, and cognitive symptoms(Jia *et al*., 2019). This process depends on multiple cellular and molecular factors, including the maturation of OPCs, the capacity of OLGs to remyelinate injured axons adequately, and molecular signals between axons and OLGs, among others (Jiang *et al*., 2023).

Tau is a microtubule-associated protein encoded by a single gene comprising 16 exons. The alternative splicing of exons 2, 3, and 10 results in six isoforms of Tau, all of them present in the adult brain (Qian and Liu, 2014). The skipping of exon 10, which encodes a microtubule-binding domain (MBD), results in the 3RTau isoform, with three MBDs, whereas its inclusion promotes the 4RTau isoform, with four MBDs. Both isoforms are present in adult brain OLGs (Villa González *et al*., 2020). 3RTau and 4RTau have a different impact on the dynamism of microtubules, the former being related to polymerization and the latter being more involved in microtubule stabilization (Qian and Liu, 2014). The relative amount of 3RTau/4RTau is indicative of the dynamics of the microtubule cytoskeleton. In the healthy adult human and rat brain, the ratio of these isoforms is 1:1. However, after induction of ischemia in rats the amount of 3RTau increases in OLGs located in the damaged area (Villa González *et al*., 2020).

Here we characterized the population of OLGs expressing high levels of 3RTau in the damaged area and their origin using a rat model of ischemia (pMCAo; permanent Middle Cerebral Artery occlusion). Of greater relevance, we demonstrate spontaneous long-term remyelination after ischemic damage, paralleled by an improvement in somatosensory reflex and motor capacities of pMCAo rats. However, using postmortem human brain samples, we ascertain that patients who died from stroke did not show the tissue response observed in the ischemic animals. These differences lead us to propose that this cellular mechanism is a self-repair brain response to long-term ischemic damage. A full understanding of this spontaneous remyelination might provide a new oligodendrocyte-based strategy to reduce the devastating effects of cerebral ischemia.

## Materials and methods

### Animals

Wistar rats aged between 2 and 3 months (200-320g) were housed in the Animal Facility of the Centro de Biología Molecular “Severo Ochoa” (CBMSO) under 12 h light/dark cycle in a temperature-controlled environment. The rats were maintained with food and water ad libitum. All the animal care protocols were conformed with national legislation (RD 53/2013) and guidelines of the European Commission for housing and care of laboratory animals. The Bio-Ethics Committee of CBMSO, the Spanish Research Council (CSIC), and the Consejería de Medioambiente de la Comunidad de Madrid (PROEX-042_5_21) approved all procedures.

We used a total number of 46 rats of both sexes (23 females and 23 males) as recommended (Lyden *et al*., 2021). Of these, 9 were used for electronic microscopy experiments, 23 for histological staining, and 14 for Western blot assays. Animals were randomly distributed into the following experimental groups: Sham, n=11 or permanent middle cerebral artery occlusion (pMCAo), n=35. The ischemic animals were distributed in three groups accordingly the number of days post-surgery named: pMCAO-2d, n=9; pMCAO5-d, n=12; and pMCAO-21d, n=14

### Permanent Middle Cerebral Artery occlusion (pMCAo)

The permanent occlusion of the middle cerebral artery (pMCAo) was performed as we described previously (Villa González *et al*., 2020). The induction of anesthesia was conducted with 3% isoflurane and the percentage was reduced at 1-1,2% for maintenance during the rest of the surgical procedure. The origin of right middle cerebral artery (MCA) was occluded by an intraluminal 4-0 Dafilon nylon suture (B. Braun, Tuttlingen, Germany) with a rounded tip coated with poly-L-Lysine (0.1% wt/vol, in deionized water; Sigma, St Louis MO). The intraluminal suture was secured with a ligature and remained in place until sacrifice 2, 5, or 21 days (2d, 5d, and 21d) post-surgery. Sham-operated animals underwent the same surgical procedures but without suture insertion. To prevent pain, all animals received skin anesthesia with lidocaine and prilocaine (25 mg/g; EMLA, Astra Zeneca).

To determine cell proliferation after pMCAo, 9 rats from the Sham, pMCAo-5d, and pMCAo-21d group received 4 intraperitoneal injections of 50 mg/kg of 5-Chloro-2′-deoxyuridine (CldU, Sigma). The injections were administered 6, 24, 48 and 72 h post-pMCAO.

### Behavioral tests

To determine the impact of pMCAo and the degree of brain function recovery long term, behavioral tests were conducted by a blind investigator at different times post-surgery.

#### The Neurological deficit score

was calculated before surgery, and at 6 h, and 1, 2, 5 and 21 days post-pMCAo using a previously described method (Villa González *et al*., 2020) based on a seven-point scale with values ranging between 0 and 6, allowing us to calculate the ischemic damage index (IDI). We excluded all ischemic rats that did not reach an IDI of 2–3 at 6 h post-pMCAo. Additionally, we tested for other neurological abnormalities, such as alterations in balance, sensorial perception, and some reflex responses (corneal, palpebral, postural correction).

#### The Beam walking test

was conducted to evaluate motor function. Rats were allowed to cross a rectangular wooden beam with a 3 cm wide and 1 m length. Operated rats were placed at one end of the beam and their ability to cross it was evaluated using a five-point scale as previously described with slight modifications (Chen *et al*., 2019). The following scoring system was used, 0-the rat could not grasp the beam and fell; 1-the rat grasped the beam and stay on it for 1 min but could not move along it; 2-the rat maintained balance but could not cross the beam; 3-the rat moved along the beam but feet slipping occurred; and 4-the rat moved freely along the beam and reached the other end without any difficulty. This test was performed in three consecutive trials 1, 2, 5 and 21 days post-pMCAO.

#### The Vibrissae-evoked forelimb-placing test

was conducted to assess somatosensory function, specifically the extension reflex of ipsilateral forelimbs. This reflex involved stimulating the vibrissae to trigger a response of placing the ipsilateral forelimb. The test was performed in five consecutive trials at 6 h, and 1, 2, 5 and 21 days post-pMCAO. The test was scored on the basis of the number of successful forelimb placements in five trials (Hua *et al*., 2002).

### Rat tissue collection

For Western blot, animals were sacrificed by exposure to CO_2_. The right parietal cortex was removed and maintained at −80°C for subsequent analysis. For histological staining, rats were deeply anesthetized using an overdose of thiobarbital (60 mg/kg) ip, followed by intracardiac perfusion with ice-cold PBS and 4% paraformaldehyde in PBS, pH=7.4 (4% PFA), as described (Villa González *et al*., 2020). Brains were post-fixed at 4°C overnight by 4% PFA immersion and cryoprotected using a sequential immersion in sucrose gradient (10% and 30%) for 48-72 hours at 4°C. After all, brains were embedded in Tissue-Tek medium (Sakura, Zoeterwoude, NL) and kept at −20°C until its cut in a cryostat (Thermo-Fisher). Coronal sections (25 μm thick) from Bregma −2 to +2 (Paxinos and Watson, 2007) were collected onto Superfrost Ultra Plus slides (Thermo Scientific, Germany) air dried at least 2 h at room temperature (RT) and stored at −20°C until use.

### Nissl method and Black Gold II staining

To quantify the ischemic area, three consecutive brain slices of each animal belonging of Bregma −2 to +2 (Paxinos and Watson, 2007) were stained using the Nissl method(Villa González *et al*., 2020). The samples from different experimental groups were stained using cresyl violet (2% in acetate buffer, Sigma). Subsequently, samples were dehydrated with sequentially increasing ethanol solutions followed by xylene and mounted with DEPEX mounting media (PanReact). The damaged area was assessed on the basis of morphological criteria previously described (Villa González *et al*., 2020), as a percentage of damaged area with respect to the total area of the hemisphere, using ImageJ software (Schindelin *et al*., 2012). The entire right hemisphere of each brain section was photographed using the “Process Manager” software tool (CellSensDimension, Olympus).

To quantify brain myelination, three consecutive slices were staining using Black Gold II (Biosensis) following the manufactureŕs instructions. Brain sections were rehydrated with distilled water for 2 min and immersed in Black Gold II at 65°C for 12 min. Slides were washed with water and then incubated with 1% sodium thiosulfate solution. Finally, the sections were dehydrated and mounted using DEPEX. The percentage of Black Gold II-positive area was measured using a custom-made macro in ImageJ software. The images were enhanced and normalized for contrast, and a color threshold was applied to eliminate the background. The results were expressed as the percentage of the Black Gold II-positive area compared to the Sham group.

The stained slides were observed under a light field microscope (Olympus BX63) and the images were captured using an Olympus DP74 camera.

### Mouse oligodendrocyte primary culture

A primary culture of OPCs was obtained from neonatal mouse pups at P2 as described (Chen *et al*., 2007). The isolated cells were plated on flasks coated with 0.1 mg/mL poly-L-lysine. OPCs were incubated in DMEM supplemented with 10% fetal bovine serum, 4.5 g/L D-Glucose, 2 nM GlutaMax, 10 ng/mL PDGF-AA (Peprotech) and 10 ng/mL of basic FGF (Peprotech). After 10 days, the culture was shaken for 1 h at 200 rpm to remove microglia. OPCs were then removed by pipetting and plated on coated-coverslips (0.1 mg/mL poly-L-lysine, 0.05 mg/mL poly-L-ornithine and 0.01 mg/mL laminin; Sigma). Differentiation of OPCs to mature OLGs was induced by incubation in DMEN/F-12 (Gibco) supplemented with 25 ng/mL insulin, 60 ng/mL progesterone, 40 ng/mL sodium selenite, 0.8 µg/mL L-Thyroxine, 100 µg/mL apo-transferrin and 20 µg/mL putrescin (Sigma) for 8 days. Cells were maintained in an incubator at 37°C with 5% CO_2_ for four days to obtain premyelinating OLGs and for 8 days to obtain matures OLGs.

### Immunofluorescence

The immunofluorescence performed in rat brain sections and mouse primary culture cells was assessed as described (Villa González *et al*., 2020). To reduced autofluorescence the samples were incubated 30 min with 0.1M glycine in PBS, pH=8,5. Subsequently, the tissue was permeabilized with 0,25% Tx-100 in PBS for 15 min and after incubated in blocking buffer containing 1% horse serum (HS), 1% BSA in PBS with 0.1% Tx-100 (PBS-Tx) for 1h at RT as described previously (Villa González *et al*., 2020). Certain primary antibodies required a previous epitope-unmasking treatment by incubation at 85°C in 10mM citrate buffer, pH=6 for 30 min. The brain samples stained with anti-BrdU were previously incubated with 2N HCl at 35°C for 30 min. The samples were incubated with the following primary antibodies overnight at 4°C, diluted in the blocking solution (1% horse serum, 1% BSA in PBS with 0.1% Tx-100): rat anti-3R-Tau 1:100 (Wako, #016-2658); mouse anti-3R-Tau 1:200 (Merck, #05-803); mouse anti-APC (clone CC1) 1:100 (Abcam, #ab16794); rabbit anti-NG2 1:200 (Millipore, #AB532); rabbit anti-Olig2 1:200 (Millipore, #AB9610); mouse anti-Olig2 1:100 (Millipore, #MABN50); rabbit anti-PDGFRa 1:200 (Abcam, #ab20349); rat anti-BrdU 1:250 (Abcam, #ab6326); and rabbit-anti-BCAS1 1:1000 (ThermoFisher, PA5-20904). Subsequently, the samples were incubated with the specific secondary antibodies (1:500; AlexaFluor, ThermoFisher) and mounted with Fluoromount-G (SouthernBiotech). Some samples were counterstained with DAPI (1:5000, Millipore) and mounted with Fluoromount-G ® (SouthernBiotech).

We used a minimum of three serial brain slices to quantify cell numbers. The images of six distinct fields from each slice were captured at 20X magnification. The number of positive cells, expressed as the number per mm^2^, was calculated using a custom-made macro with Fiji software. In brief, the image contrast was enhanced and then a threshold was established to eliminate the background. We used the “Analyze particles” tool to quantify Olig2^+^/CC1^+^ cells with a size=200-Infinity and circularity= 0.08-1. To quantify Olig2^+^/NG2^+^, PDGFRα^+,^ and BCAS1^+^/Olig2^+^ cells, we used a “cell counting” tool. The images were obtained with the 405 confocal LSM710 vertical line.

### Human brain samples and immunohistochemistry

The brain samples and data from patients included in this study (Table 1; 5 stroke patients and 5 controls) were provided by the Biobank HUB-ICO-IDIBELL (PT20/00171), integrated into the ISCIII Biobanks and Biomodels Platform, following the guidelines of Spanish legislation (RD 1716/2011) and with approval of the local ethics committees (Comite de ética de investigación CEI-UAM and Comité Científico Externo CCE-Biobanc HUB-ICO-IDIBELL). Samples were anonymized and processed in a blinded manner. Brain tissue was consistently sampled from similar regions of the cerebral cortex. Samples from controls were collected from the occipital cortex. Samples from stroke patients were selected on the basis of the extent of damage to the anterior cingulate, frontal, occipital, or parietal cortex. Post-mortem delays were ranged between 3 to 14 h for all the samples. All the brain tissue samples were fixed with 10% formalin and embedded in paraffin using a conventional method. The sections were cut at 10 µm thickness and Nissl stained to determine the damaged area (see “Nissl method and Black Gold II staining” section). Additional slices were used for the immunohistochemical studies using Olig2, 3RTau, and MBP antibodies.

**TABLE 1:** Human samples and brain areas included in this study.

Summary of the patients from whom cortical brain samples were used in this studio. M: male; F: female; C: control, S: stroke.
| Sample code | Age (years) | Sex | Diagnosis | Death Cause | Postmortem Delay hours |
| --- | --- | --- | --- | --- | --- |
| C1 | 66 | M | C | Cardiac failure | 3 |
| C2 | 33 | M | C | Hypovolemic shock | 10 |
| C3 | 56 | M | C | Liver failure | 10 |
| C4 | 57 | F | C | Respiratory failure | 15 |
| C5 | 53 | F | C | Infectious disease | 5 |
| S1 | 54 | M | S | STROKE: infarction, middle cerebral artery | 3 |
| S2 | 76 | M | S | STROKE: infarction, middle cerebral artery | 4 |
| S3 | 73 | F | S | STROKE: infarction, middle cerebral artery | 16 |
| S4 | 65 | M | S | STROKE: infarction, middle cerebral artery | 11 |
| S5 | 70 | F | S | STROKE: infarction, middle cerebral artery | 14 |

Samples were deparaffined by heating them to 62°C for 30 min and washed in xylol 2 times. Then, were rehydrated with decreasing ethanol concentrations. Antigen retrieval was applied by heat-induced epitope retrieval in 10 mM Tris-EDTA buffer, pH=9.0, or 10 mM citrate buffer, pH=6.0, depending on the primary antibody, for 13 min. Samples were then blocked in a buffer containing 1% HS in PBS-Tx, pH=7.4 for 1 h, followed by incubation with the appropriate primary antibodies diluted in blocking solution: rat anti-3R-Tau 1:100, rabbit anti-Olig2 1:200, and anti-MBP 1:200 at 4°C. Brain slices were incubated with 1% H_2_O_2_ to block endogenous peroxidase and then with the appropriate biotinylated secondary antibody using the VECTASTAIN-ABC kit (Vector Laboratories Inc.), following the manufactureŕs instructions. Staining was performed by incubation with FAST 3,3′-diaminobenzidine tablets (Sigma). Finally, brain sections were dehydrated using ethanol gradient, cleared with xylol and mounted using DEPEX.

To determine the number of Olig2^+^ and 3R-Tau^+^ cells, we used a custom-made macro in Fiji software (see “Immunofluorescence” section) using the specific parameters of size=150–1000 and circularity= 0.4–1. The number of positive cells was expressed as the mean of the total cells counted in three consecutive fields of the subcortical white matter.

Stained slides were observed under light field microscope (Olympus BX63) and the images were captured using an Olympus DP74 camera.

### Transmission electronic microscopy (TEM)

Samples for TEM analysis were obtained from Sham, pMCAo-5d, and pMCAO-21d rats (n=3 animals per experimental group) transcardially perfused with 2% glutaraldehyde/4% PFA in 0.1M phosphate buffer. The brains were removed and fixed overnight at 4°C by immersion. The ipsilateral hemisphere was cut into 200 µm sections using a vibratome (Leica VT1200) and stored in PBS until use. Subsequently, a post-fixation step with 1% osmium tetroxide and 0.8% potassium ferricyanide was performed for 1 h at 4°C. Next, the samples were incubated with 0.15% tanic acid for 1 min at RT and subsequently with 2% uranyl acetate for 1 h in the dark. They then underwent a solvent-based dehydration step, followed by embedding in TAAB 812 epoxy resin (TAAB Laboratories, Berkshire, UK) before ultra-fine sectioning and incubated with a heavy metal stain. Ultra-fine sections of 70 nm were cut with an Ultracut E ultramicrotome (Leica Systems, Wetzlar, Germany) and mounted on Formvar-carbon-coated Cu/Pd slot grids. The heavy metal staining was performed with uranyl acetate and lead citrate.

Samples were viewed under a JEM-1400 Flash electron microscope (JEOL Ltd., Tokyo, Japan) at 100 kV, and a OneView 4 k × 4 k CMOS camera (Gatan, Pleasanton, CA) was used to acquire images. The micrographs were taken and assessed by a blinded researcher.

Myelinated and unmyelinated axons were quantified by counting the total number of axons in three fields of view per mm^2^ of each animal (minimum of 800 axons per sample). Data were represented as a percentage of myelinated or unmyelinated axons of the total axons counted for each rat. The diameter of each axon was determined using the ImageJ measurement tool (minimum of 300 axons per animal) and classified as large (≥1 µm), medium (between 0.7 and 0.99 µm), and small (≤0.69 µm). The g-ratio was calculated by dividing the inner axonal diameter by the outer axonal caliber of 100–200 axons per animal in three steps, as previously described (Christina *et al*., 2020).

### Western blot

Western blot of homogenates of the right parietal cortex of rats was performed as described (Villa González *et al*., 2020). Right parietal cortex of rats was homogenized in a buffer containing: 50 mM Tris pH = 8, 150 mM NaCl, 1% Triton X-100, 5 mM EDTA, protease inhibitor cocktail (Sigma-Aldrich) and phosphatase inhibitors (2 nM okadaic acid and 1 mM Na3VO4) for 30 min at 4°C. Samples were then centrifuged at 10,000g for 5 min and the supernatants were collected. The protein concentration of each sample was determined using the DC Protein Assay kit (BioRad, Hercules, CA). The proteins were resolved by SDS-PAGE using a Mini-Protean system (BioRad). The total proteins (40 μg) were loaded in a buffer containing: 0.063 mM Tris pH = 6.8, 10% SDS, 25% glycerol, 5% β-mercaptoethanol, and 0.5% bromophenol blue. Samples were heated at 95°C for 5 min before loading. After electrophoresis, the proteins were transferred into nitrocellulose membranes (Merck-Millipore, Germany) for 2h at 300 mV using an electrophoretic transfer system (BioRad). The membranes were then blocked for 1 h with 5% of skimmed milk in PBS containing 0.1% Twen-20 (PBS-T) and incubated overnight at 4°C with the corresponding primary antibodies diluted in blocking buffer: mouse anti-β-Actin 1:2000 (Sigma, #A544); mouse anti-3R-Tau 1:1000; rabbit anti-PLP 1:1000 (Abcam, #ab254363); mouse anti-MOG 1:1000 (Sigma, #MAb91087); rabbit anti-MAG 1:1000 (Proteintech, #14386); and rat anti-MBP 1:500 (Millipore, #MAB386). Membranes were rinsed with PBS-T and incubated with the subsequent secondary antibody, namely anti-rabbit IgG-HRP (Sigma), anti-mouse IgG-HRP (Sigma), or anti-rat IgG-HRP (Sigma). Immunoreactivity was detected by the Clarity Western ECL substrate (BioRad). The relative density of the bands was calculated from their optical density using Fiji software. The values were normalized to actin levels detected in the same membrane and compared with control values (Sham).

### Statistical analysis

All statistical tests were performed with GraphPad Prism 8. The data were expressed as mean ± SEM. A Shapiro–Wilk test was used to check for normal distribution of each data set. The parameters named g-ratio and axon diameter are represented as individual values. When comparing more than two experimental groups, a two-way analysis of variance (ANOVA) followed by multiple comparisons with Bonferroni correction was performed. When two groups were compared, an unpaired t-test was used. For the statistical analysis of g-ratio and axon diameter, we used the Kruskal-Wallis test. All statistical tests were performed in a two-tail manner.

## Results

### Long-term spontaneous improvement of neurological and sensorimotor functions 21 days after pMCAo

Quantification of the damaged area, using Nissl staining (Fig. 1A-1C), revealed a progressive increase following pMCAo, reaching 65%±5 (p≤0.001) of the ipsilateral hemisphere 21 days post-surgery. Neurobehavioral tests conducted 6 h post-pMCAo showed an IDI of 2.15, indicating the successful induction of ischemia. Five days after surgery, the IDI score increased, reaching those of Sham rats at 21 days post-surgery. To ascertain whether this improvement in the IDI score was reflected by enhanced sensorimotor function associated with the parietal cortex, the region of the brain most affected by ischemia, we conducted two additional tests, namely the beam-walking and vibrissae-evoked forelimb-placing tests. Both revealed a significant reduction in the score in the short term. However, 5 days after pMCAo, the scores of the animals on the beam-walking test that were similar to those of Sham counterparts, whereas the tactile reflex was recovered at 21 days.

**Figure 1:**
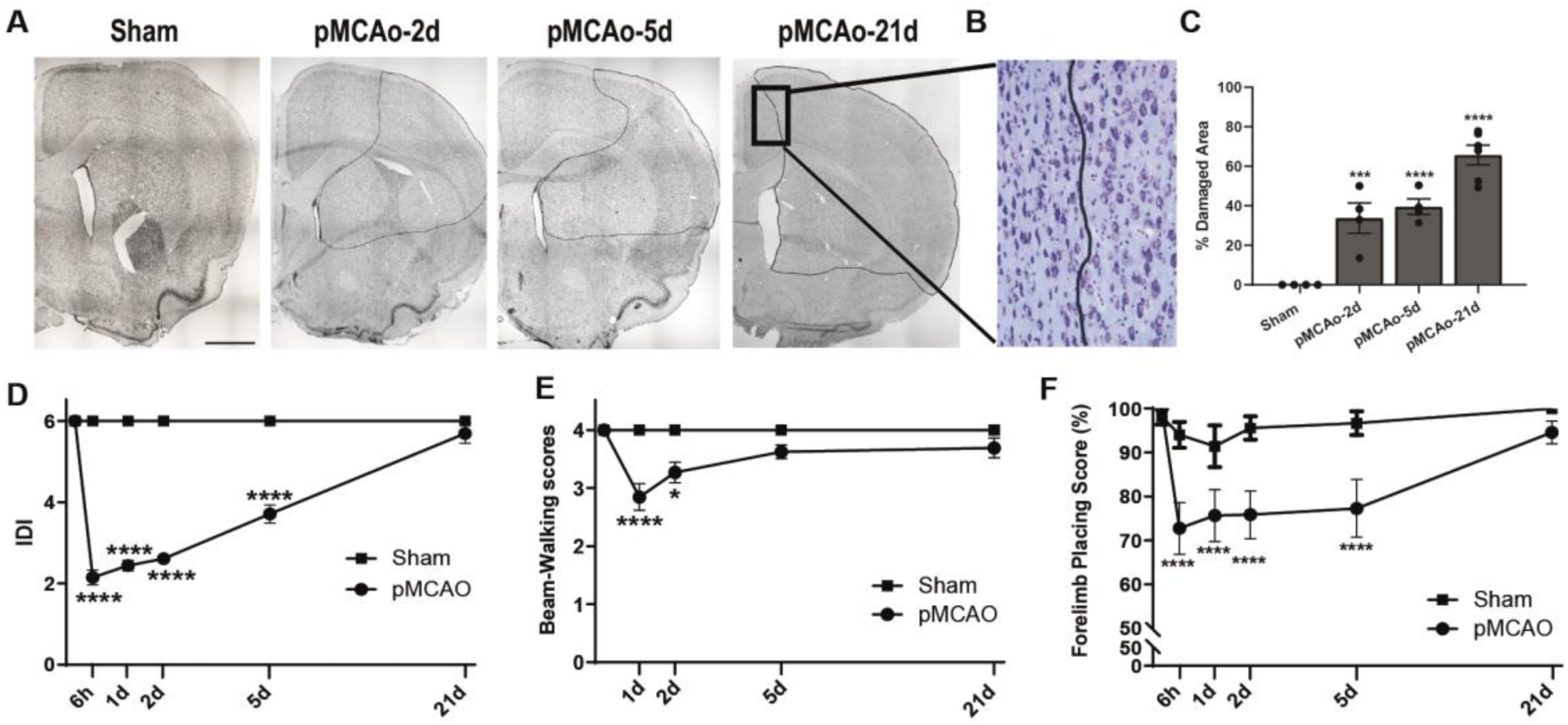
Temporal evolution of ischemic damage after pMCAo. **(A)** Representative montages of low magnification of a brain hemisphere of Nissl-stained coronal brain slides (25 µm) of control (Sham) or ischemic animals (pMCAo) 2, 5 and 21d after ischemia induction. The solid line indicates the limit of damaged area, scale bar: 200µm. **(B)** Magnification of pMCAo-21d Nissl-stained image shows damaged (right) and healthy tissue (left). **(C)** The bar-graphs represent the percentage of damaged area respect to the total of the hemisphere measured in three consecutives coronal brain sections for each rat using coronal slices of pMCAo rats at 2, 5 and 21d post-ischemia. Bars represent the mean ±SEM (n=5-6). Unpaired t-test *p<0.05, **p< 0.01, ***p<0.001 ****p<0.0001. **(D)** The ischemic damage index (IDI) measured using a seven-point scale by a standard neurological test: 0 corresponds to the lowest and 6 to the highest degree of neurological score. **(E)** Beam-Walking test measured using a four-point scale: 1 corresponds to the lowest and 4 to the highest. **(F)** Forelimb-placing score assessed as percentage of extension of the impaired forelimb of five trials. All represented data correspond to the means ±SEM of pMCAo (n=27) or Sham (n=11). Two-way ANOVA; *p<0.05,**p< 0.01, ***p<0.001 ****p<0.0001.

### Temporal dynamics of OPC and mature OLG populations following permanent ischemia

To characterize the temporal response of OLG populations to ischemic injury, we assessed the density of OPCs, and pre-myelinating and mature OLGs in six fields of damaged cortical areas (Supplementary Fig. 1B). OPCs were quantified using two specific markers, namely NG2 and PDGFRα. Given that NG2^+^ expression can also be observed in other cellular lineages(He *et al*., 2023; Huang *et al*., 2020; Kirdajova *et al*., 2021), we considered only cells double positive for NG2 and Olig2 (NG2^+^/Olig2^+^). The quantification of mature OLGs was calculated using immunofluorescence with the anti-APC (clone CC1) antibody, which specifically recognizes a protein upregulated in myelinating OLGs(Frazier *et al*., 2023). Since it has been reported that this antibody can also be detected in a subset of astrocytes(Bin *et al*., 2016), we included only CC1^+^/Olig2^+^ cells. The number of pre-myelinating OLGs was determined by counting BCAS1^+^/Olig2^+^ cells(Jiang *et al*., 2023).

Two days after pMCAo, the density of OPCs increased significantly compared to the Sham group, reaching 379 cells/mm^2^ (p≤0.05) (Fig. 2A and 2B). However, at 5 and 21 days post-mMCAo, the OPC population exhibited a lower density (100 cells/mm^2^ at 5 days and 150 cells/mm^2^ at 21 days) compared to the Sham group (240 cells/mm^2^; p≤0.05). These findings were further supported by analysis of PDFRα^+^ cells, which showed the same tendency (Supplementary Fig. 1A). BCAS1^+^/Olig2^+^ cells showed a significant increment with respect to Sham animals at 5 days post-pMCAo (366±5.2 versus 132±21 cells/mm^2^; p≤0.01) (Fig. 2A and 2C). The density of CC1^+^/Olig2^+^ cells (Fig. 2A and 2D) maintained Sham values at 2 and 21 days. However, at 5 days post-pMCAo, this population fell below that of the Sham group (1428±155 cells/mm^2^; p≤0.05). The analysis of the temporal evolution of OLG populations using the percentage of variation with respect to Sham animals (Fig. 2E) summarizes these dynamics.

**Figure 2:**
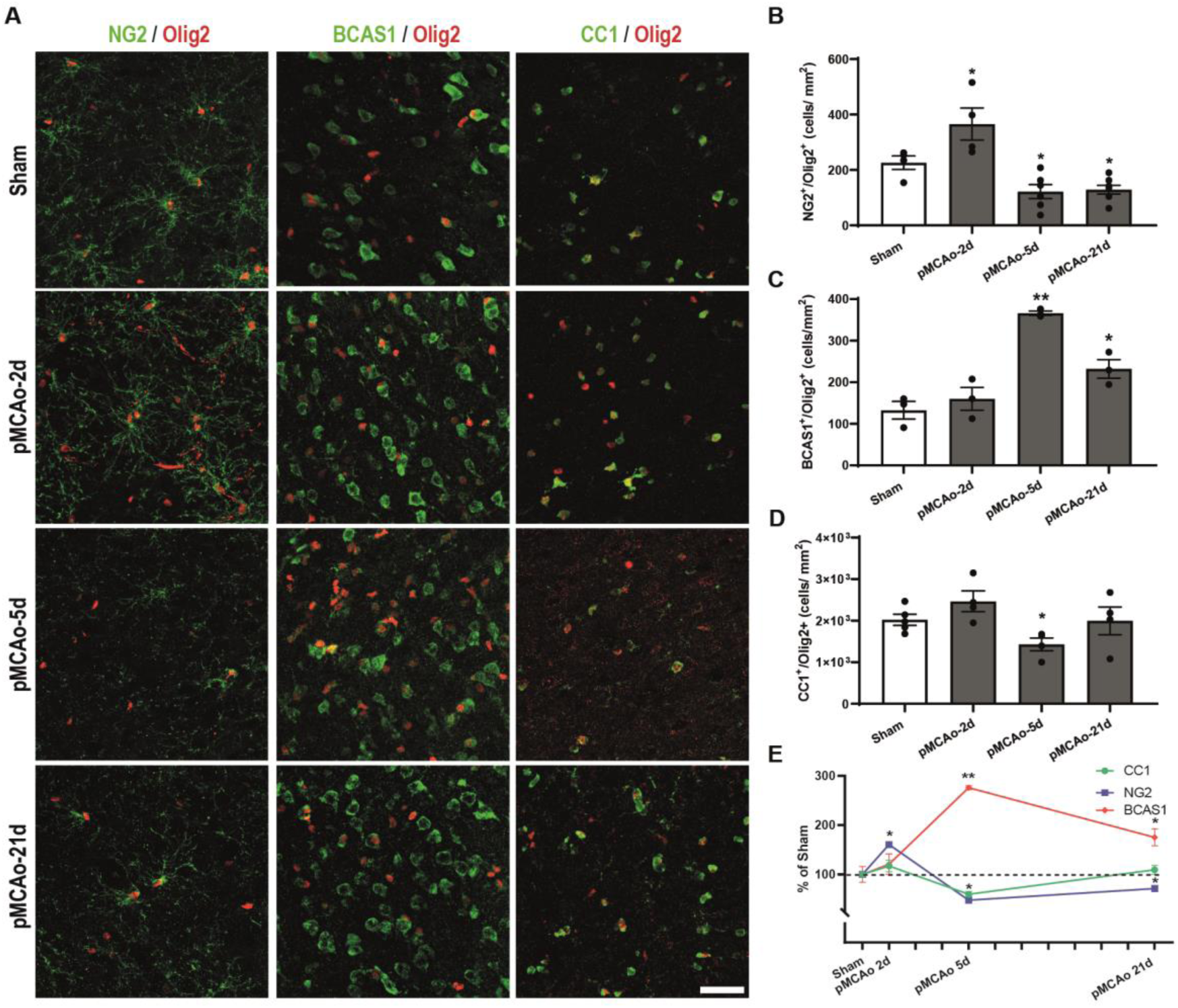
Analysis of temporal evolution of oligodendrocyte subpopulations in damaged area after cerebral ischemia. **(A)** Representative confocal microphotographs corresponding to parietal cortex of coronal brain sections (25µm) of Sham and pMCAO rats (2-, 5- and 21 days after ischemia induction), double-stained with Olig2 (red) and NG2 (left column), BCAS1 (middle column), CC1 (right column) in green. Scale bar: 40 µm. **(B)** Histograms show the density of NG2^+^/Olig2^+^, **(C)** BCAS1^+^/Olig2^+^ and **(D)** CC1^+^/Olig2^+^ cells per mm^2^ of Sham and pMCAo animals. Bars represent the mean ± SEM (n=5-6). Unpaired t-test; *p<0.05, **p<0.01. **(E)** Linear representation of the percentage of variation of NG2^+^/Olig2^+^, BCAS1^+^/Olig2^+^ and CC1^+^/Olig2^+^ compared to Sham group. Each point shows the mean ± SEM.

### pMCAo-induced increase in 3RTau^+^ OLGs in the damaged area is independent of oligodendrogenesis

Previous results demonstrated an increment in the number of 3RTau^+^/Olig2^+^ cells in the damaged area after cerebral ischemia(Villa González *et al*., 2020). To characterize this OLG population, we used immunofluorescence assays with specific antibodies to OPCs, and pre-myelinating and mature OLGs combined with anti-3RTau. No PDGFRα^+^/3RTau^+^ cells were detected in any experimental group (Fig. 3A). However, we found BCAS1^+^/3RTau^+^ and CC1^+^/3RTau^+^ cells at all the time points studied, their colocalization being confirmed by orthogonal view (Fig. 3B and 3C). These results were confirmed by immunofluorescence in mouse primary culture cells at 4 and 8 DIV (Fig. 3H and 3I). The percentage of cells expressing 3RTau^+^ reached almost 60% of total BCAS1^+^ population in Sham and pMCAo-2d groups. However, 5 and 21 after pMCAo, the percentage of 3RTau^+^ cells increased by over 80% (p≤0.01 and p≤0.05 respectively) (Fig. 3E). A similar analysis of CC1^+^ cells (Fig. 3F) revealed a significant increase 2 days after damage (p≤0.05). Using Western blot, we confirmed this increment in 3RTau levels in the parietal cortex, where it peaked at 151% (p≤0.01) in the pMCAo-5d group (Fig. 3D, 3G), coinciding with the highest density of pre-myelinating OLGs.

**Figure 3.**
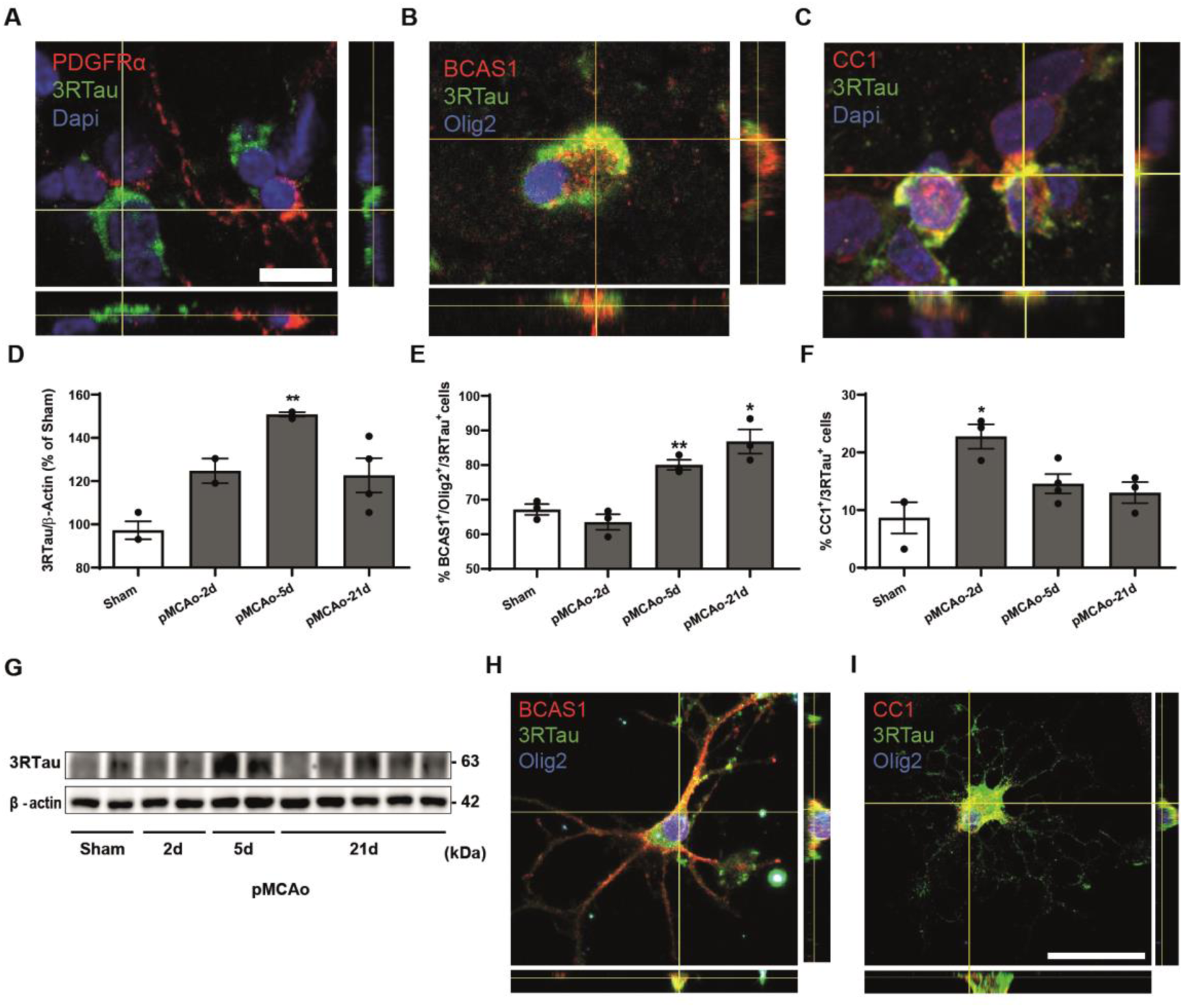
Identification of oligodendrocyte subpopulations expressing 3RTau+ in damaged area. **(A-C)** Confocal orthogonal view representative of a triple immunofluorescence of PDGFRα (red), 3RTau (green) and DAPI (blue); BCAS1 (red), 3RTau (green) and Olig2 (blue); CC1 (red), 3RTau (green) and Dapi (blue). All the images correspond to the parietal cortex of pMCAo-2d group. Scale bar: 5 µm. **(D)** Histogram shows the abundance of 3RTau normalized with β-actin in homogenates of parietal cortex of pMCAo and Sham rats, expressed as a percentage of values of Sham. Each bar represents the mean ±SEM (n =14). Student’s t-test; p-values: **p<0.01. **(E, F)** Bar-graph representing the percentage of cells triple positives BCAS1^+^/Olig2^+^/3RTau^+^respect total BCAS1^+^/Olig2^+^(E); or double positives CC1^+^/3RTau^+^ respect total CC1^+^/Olig2^+^ cells (F) of Sham and pMCAo animals (n=3 of each experimental group). Each bar shows the mean ±SEM. Unpaired t-test: *p<0.05, **p<0.01. **(G)** Representative western blot of 3RTau protein levels in ischemic area (parietal cortex). **(H, I)** Representative confocal orthogonal view of 4 (H) and 8 (I) days *in vitro* mouse oligodendrocyte primary culture stained with BCAS1 (red), 3RTau (green) and Olig2 (blue) or CC1 (red), 3RTau (green) and Olig2 (blue). Scale bar: 40 µm.

To ascertain the origin of the OLG colonizers of the ischemic area, we intraperitoneally injected the rats with CldU, a cell-division marker, after the surgical procedure. We analyzed the lateral ventricle as a source of newly generated OLGs (Fig. 4A), and the parietal cortex (damaged area; Fig. 4C). Our results indicated an increase in CldU^+^/Olig2^+^ cells in the lateral ventricle (Fig. 4A and 4B) at 5 days post-pMCAo, and a decrease at 21 days, at which time their location shifted towards more lateral brain regions, below the corpus callosum (data not shown). However, we did not detect any CldU^+^/Olig2^+^ cells in the damaged cortex (Fig. 4C and D).

**Figure 4.**
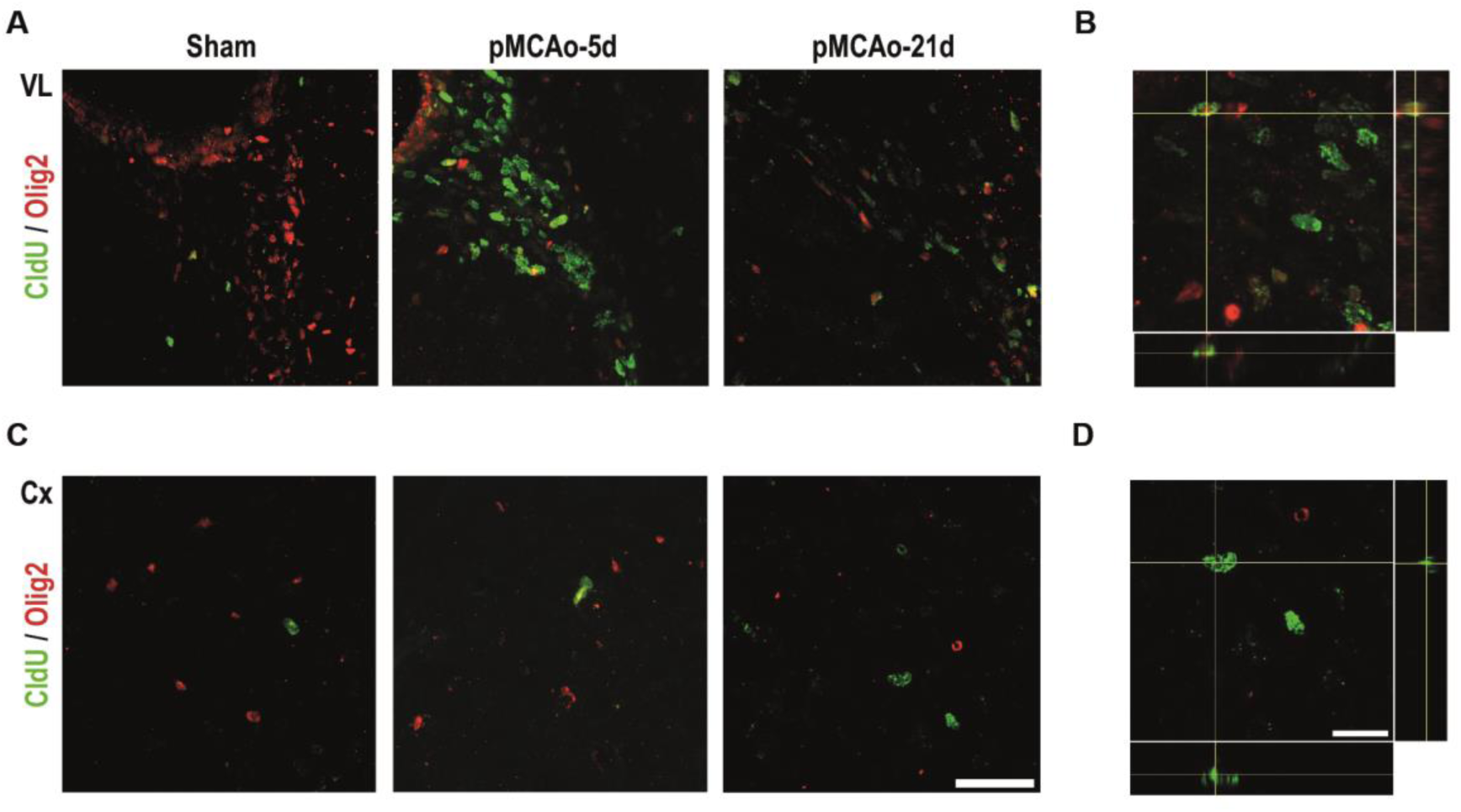
Oligodendrogenesis is exclusive of subventricular zone of lateral ventricle following ischemia. **(A and C)** Representative confocal images of double immunofluorescence with Olig2 (red) and CldU (green) of Sham and pMCAO animals of lateral ventricle (A) and parietal cortex (C). Scale bar: 40 μm. **(B and D)** Orthogonal views of double positive cells Olig2^+^ (red) and CldU^+^ (green) of lateral ventricle of pMCAo-5d rats (B) and CldU^+^ cell (green) in the cerebral cortex of pMCAo-21d animals. Scale bar: 20µm

### Long-term spontaneous remyelination after pMCAo induction

To determine whether remyelination occurs we selected two different fields in each brain slice stained with Black-Gold II (see Fig. 5B). The percentage of Black-Gold II-positive area in the pMCAo-2d and pMCAo-5d groups decreased compared to the Sham group (66%; p≤0.05 and 52%; p≤0.001, respectively), thereby indicating demyelination after ischemia (Fig. 5A, 5C). However, the Black-Gold II staining did not disclose any significant difference between the Sham and pMCAo-21d groups, thus indicating remyelination (Fig. 5A and 5C).

**Figure 5.**
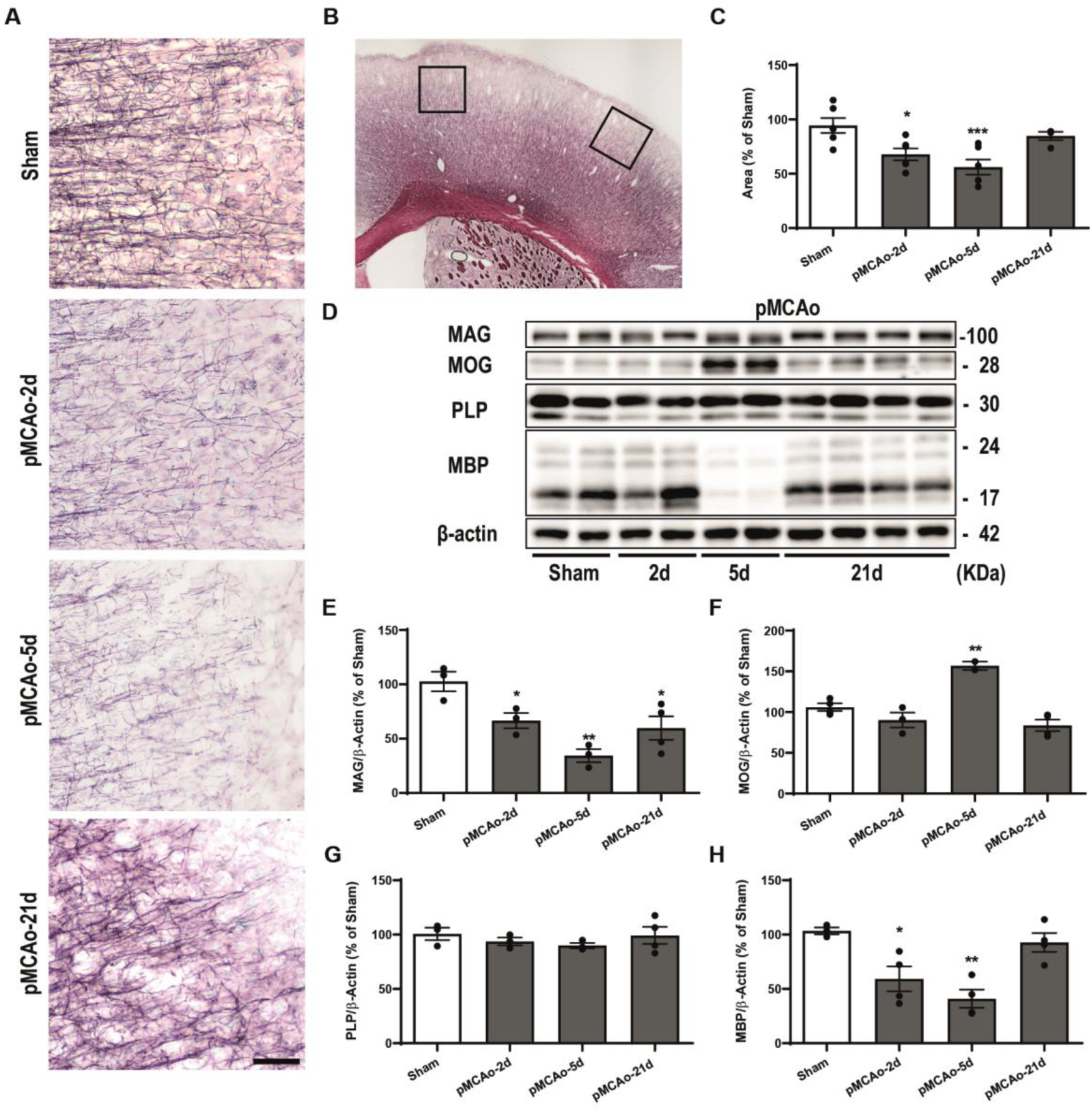
Analysis of myelin progress after pMCAo induction. **(A)** Representative Black Gold II images of the parietal cortex from Sham and pMCAO animals. Scale bar: 40 μm **(B)** Photograph of the right parietal cortex of a Black Gold II-stained coronal section. The squares indicate the fields used in this analysis. **(C)** Graphical representation of the percentage of positive Black Gold II-stained area of Sham and pMCAo groups (n=5 of each experimental group) **(D)** Representative western blots of MAG, MOG, PLP, MBP and β-actin that show the levels of these proteins in homogenates of parietal cortex of Sham and pMCAo animals. **(E-H)** Histograms show the abundance of MAG (E), MOG (F), PLP (G) and MBP (H) normalized with β-actin expressed as a percentage of values compared to Sham. Each bar represents the mean ±SEM (n =14). Unpaired Student’s t-test; *p<0.05, **p<0.01, ***p<0.001 ****p<0.0001.

We analyzed the temporal evolution of some myelin-associated proteins, namely myelin-associated glycoprotein (MAG), myelin oligodendrocyte glycoprotein (MOG), proteolipid protein (PLP), and myelin basic protein (MBP) (Fig. 5D). The levels of MAG dropped to 34±6% of those of Sham animals at 5 days post-pMCAo (Fig. 5D and 5E) (p≤0.01) and partially recovered at 21 days (59±12 respect to Sham; p≤0.05). MBP levels were drastically reduced at 2 and 5 days post-pMCAo (59%±13 and 33%±9.6 respect to Sham; p≤0.0001) and recovered at 21 days (Fig. 5D and 5H). MOG levels (Fig. 5D and 5F) showed a significant increase at 5 days post-pMCAo (157%±4.2 of Sham; p≤0.0001). However, the levels of PLP did not show significant variations with respect to Sham animals at any of the times analyzed (Fig. 5D and 5G).

TEM confirmed our previous results, revealing marked demyelination at 5 days post-pMCAo and spontaneous remyelination at 21 days (Fig. 6A). We considered axons presenting compact myelin as “myelinated” (Fig. 6G and 6H). The round axons with myelin aberrations as vacuoles (Fig. 6I) and round axons with more than 25% of the total perimeter of uncompacted myelin layers were considered “unmyelinated” (Fig. 6J). The percentage of myelinated axons in the pMCAo-5d group was significantly reduced (12±6% of total counted axons; p≤0.05) but reached the Sham values on day 21 (Fig. 6B). An analysis of the percentage of myelinated axons on the basis of their caliber revealed a marked reduction of myelinated axons of all sizes in the pMCAo-5d group, this decrease being more notable for the largest ones. At 21 days post-pMCAo, the percentage of myelinated axons almost reached that of the Sham animals, with no significant differences in any axon size (Fig. 6C). However, analysis of the average diameter of myelinated axons (Fig. 6D) showed a progressive and significant decrease in axon caliber after ischemia induction (Sham: 0.88±0,07µm; 5d-pMCAo 0.70±0.09 µm and 21d-pMCAo 0.64±07µm; p<0.0001).

**Figure 6.**
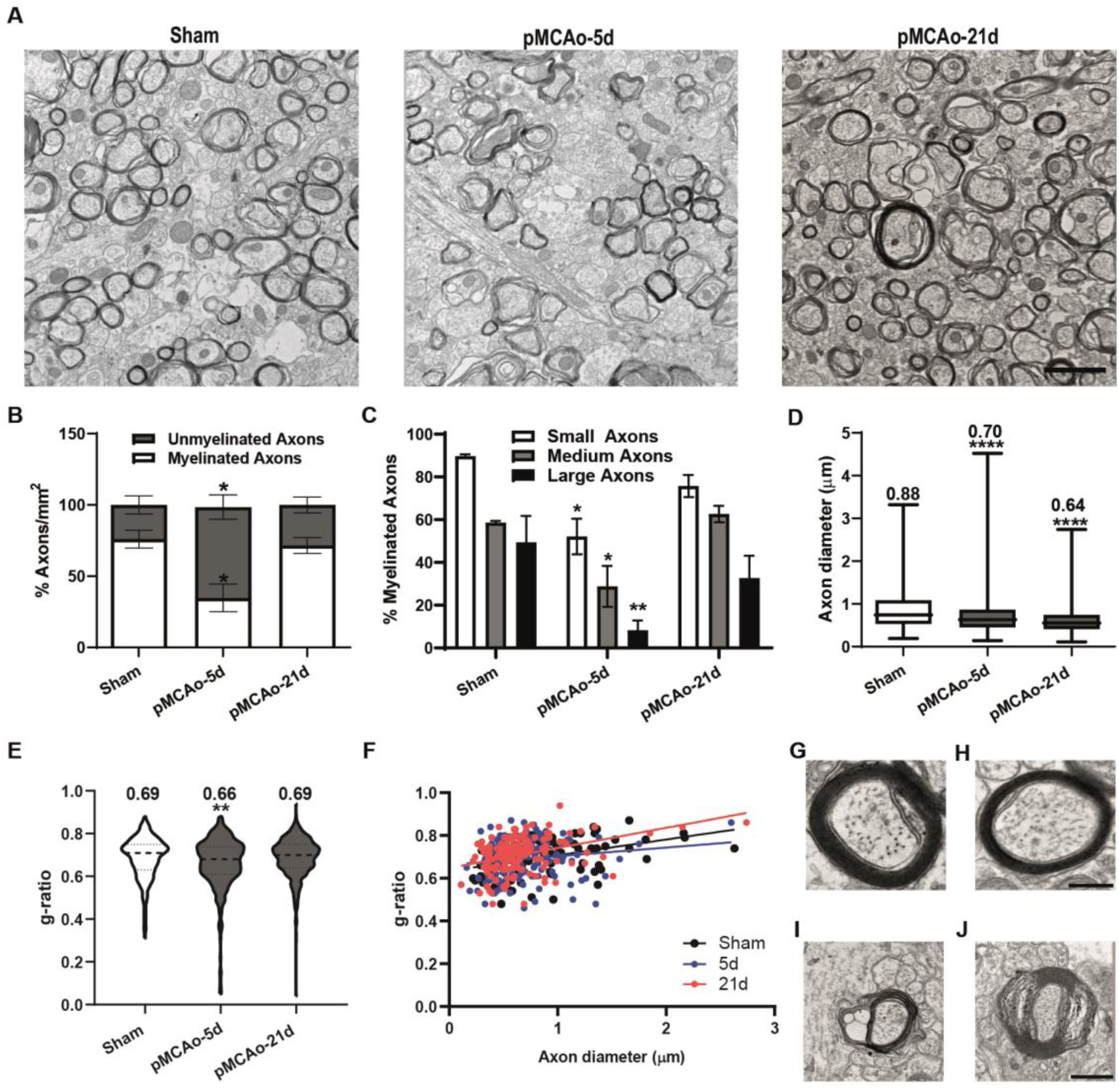
Ultrastructural analysis of myelin after brain ischemia. **(A)** Representative electron transmission microscopy images of right parietal cortex from Sham, pMCAo-5d and pMCAO-21d groups. Scale bar: 2 μm. **(B)** Histograms show the percentage of myelinated and unmyelinated axons of a minimum of 800 total axons counted per individual of each experimental group. Two-way ANOVA followed by Bonferroni multiple comparisons. **(C)** Graphical representation of the size-based distribution of myelinated axons of each experimental group. The axons were grouped according with their size in large (≥1 µm), medium (≥0.7≤0.99 µm) and small (≤0. 69 µm). The bar represents the mean ±SEM of the percentage of large, medium or small axons of all myelinated axons counted. **(D)** The box plot shows the mean ±SEM of the diameter of myelinated axons of each experimental group: Sham, pMCAo-5d and pMCAo-21d. **(E)** Violin diagram representes the average of g-ratio of all the myelinated axons counted of Sham and pMCAO rats. **(F)** The scatter plots fitted to a linear function shows g-ratio and diameter relationships of Sham (black), pMCAo-5d (blue) and pMCAo-21d (red). **(G-H)** Representative image of a myelinated axon of Sham (G) and pMCAo-21d groups (H). Scale bar: 250nm. **(I-J)** Representative images of unmyelinated axons with aberrations as vacuoles (I) or delaminated myelin “onion like” (J). Scale bar: 500nm. Two-way ANOVA followed by Bonferroni multiple comparisons was used to compare the percentage of axons/mm^2^ and percentage of myelinated axons. To compare the average axons diameter and the average g-ratio, an unpaired t-test was used (n=3), p values: *p<0.05, **p<0.01, ***p<0.001 ****p<0.0001.

The average g-ratio, a measurement of myelin sheath thickness proportional to the diameter of myelinated axons, is a commonly used indicator of a structural trait that affects nerve impulse conduction (Fig. 6E). At 5 days post-pMCAo, the mean g-ratio decreased with respect to the Sham group (0.66±0,01% versus 0.69±0.02; p≤0.0001), but at 21 days this parameter recovered Sham values. Analysis of the relationship between the g-ratio and axon diameter, using scatter plot fit with a linear function, showed that large axons were the most affected, with a lower g-ratio in the pMCAo-5d group (Fig. 6F) and a higher one in the pMCAo-21d group compared to Sham animals (Fig. 6F, 6G, 6H).

### 3RTau^+^ cells are reduced in the subcortical white matter of stroke patients

Postmortem brain samples from stroke patients stained with Nissl showed clear tissular damage (Fig. 7A) with a lower cell density, a paler cresyl violet stain, and shrunken cell bodies compared to controls. Furthermore, using specific markers for OLGs (Fig. 7B) and 3RTau (Fig. 7C), we observed a dramatic reduction in the number of Olig2^+^ (104 ±11 cells versus 221 ±9; p≤ 0.0001) (Fig. 7B and 7F) and 3RTau^+^ cells (24±6 cells versus 74±3; p ≤ 0.01) (Fig. 7C and 7G) in subcortical white matter. MBP staining (Fig. 7D) revealed aberrantly condensed and less complex myelin in the same area in the stroke group. To quantify this, we used the coherency of myelinated axons parameter, considered an inverse measure of the complexity of white matter microstructure, using the Fiji tool “OrientationJ measure”(van Tilborg *et al*., 2017). Compared to control samples, those from stroke patients showed a significant increase in coherency (0.15±0.0; p<0.01) (Fig. 7H), thereby indicating a reduction in the microstructural complexity of myelin.

**Figure 7.**
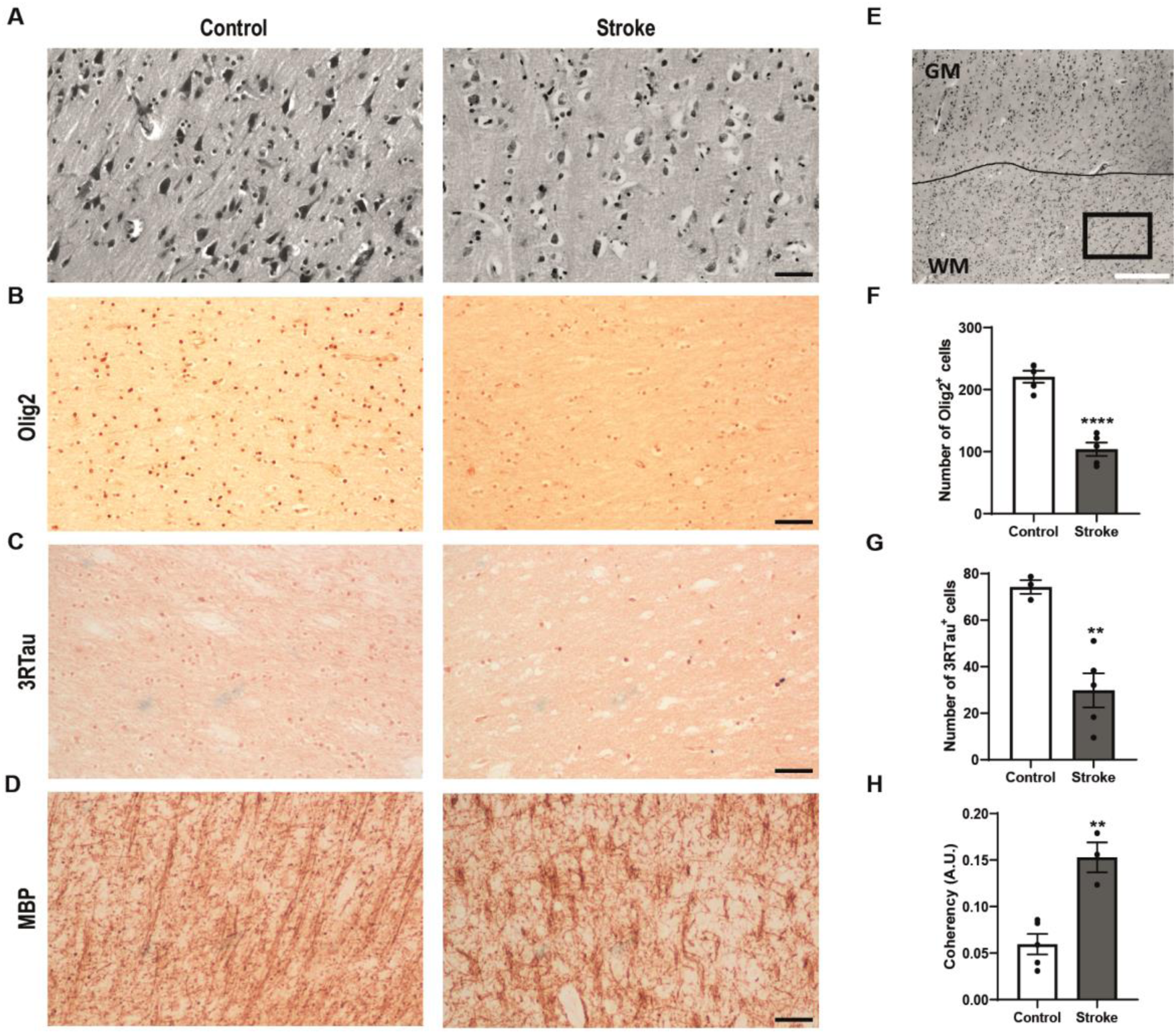
Postmortem human brain samples from stroke donors present lower levels of 3RTau^+^ cells in subcortical white matter than control. **(A)** Representative Nissl-stained images of cerebral cortex slices from control and stroke donors. **(B-D)** Selected images of subcortical white-matter stained with anti-Olig2 (B), anti-3RTau (C) and anti-MBP (D) antibodies of control (n=5) and stroke (n=5) donors. Scale bar: 200 μm. **(E)** Nissl-stained coronal section of cerebral cortex of control donor. The line separates gray matter (GM) from white matter (WM). Square represents the selected area for Olig2, 3RTau and MBP analysis. **(F-H)** Bar graphs shows the mean ±SEM of the total number of Olig2^+^, F, 3RTau^+^ cells, G, and coherency of myelinated axons, H, as a parameter to analyzed myelin complexity. Unpaired t test: *p<0.05, **p<0.01, ***p<0.001 ****p<0.0001.

## Discussion

Ischemic stroke is one of the foremost public health problems worldwide and it carries a high economic burden (Campbell *et al*., 2019). Most studies performed in this field have focused on the acute phase of the disease in order to reduce its impact and have overlooked chronic phases. Some stroke patients and animal models spontaneously experience a significant reduction of their symptoms and even long-term recovery of some brain functions, and such cases are still not fully understood (Jia *et al*., 2019; Villa González *et al*., 2020). This spontaneous improvement points to the functional restoration of some cerebral circuits and thus reflects the ability of the injured brain to trigger a self-repair process to reduce ischemic damage in some cases.

Myelin increases the processing speed of neural circuits and provides trophic support to axons, thereby reducing neuronal damage (Hernández *et al*., 2021). The “classical view” of permanent static myelin in the healthy adult brain is now shifting towards a more dynamic vision of this structure(Chapman and Hill, 2020). Recent observations strongly suggest that myelination serves as a form of adult plasticity by fine-tuning cerebral circuits (Hughes *et al*., 2018; Liu *et al*., 2012). Some authors have proposed that this plasticity, through remyelination, serves as a potential brain self-repair mechanism after damage (Shi *et al*., 2015). A greater understanding of this process is key to identifying the failure of regenerative mechanisms in some patients. Hence, there is increasing interest in unraveling the OLG response to brain damage.

Analysis of the temporal evolution of OLG populations in the damaged region of the brain showed an increase in the density of OPCs 2 days after ischemia, as previously described (Bonfanti *et al*., 2017; Chu *et al*., 2012). Named polydendrocytes, NG2^+^ cells are considered a fourth type of glial cell in the adult CNS and they have differentiation potential under physiological and ischemic conditions (Bonfanti *et al*., 2017). After brain ischemia, some NG2^+^ cells express markers of other lineages (He *et al*., 2023; Huang *et al*., 2020; Kirdajova *et al*., 2021), thus suggesting that they fail to differentiate into mature OLGs. Our results using two different approaches allowed us to evaluate the temporal evolution of OPCs, avoiding other possible cellular lineages, and strongly support colonization of the damaged area by OPCs, an observation consistent with other authors (Chu *et al*., 2012). Pre-myelinating OLGs (BCAS1^+^/Olig2^+^) showed a significant increase in density 5 days after damage, when OPCs and mature OLGs exhibited the lowest densities. The special sensitivity of OLGs to ischemia (Dewar *et al*., 2003; Hernández *et al*., 2021) may explain the results observed 5 days post-pMCAo. Meanwhile, CC1^+^/Olig2^+^ cells returned to Sham levels 21 days after damage. These results suggest that OPCs differentiate into mature OLGs, in contrast to some reports that indicated proliferation and migration of OPCs that fail to differentiate to mature-myelinating OLGs to the damaged area after ischemia (Bonfanti *et al*., 2017; Chu *et al*., 2012; He *et al*., 2023; Song *et al*., 2017). Similarly, Bonfanti et al. (Bonfanti *et al*., 2017), using transgenic GPR17iCreER(T2):CAG-eGFP mice under pMCAo induction, described that some OPCs recruited to the ischemic area acquire mature OLG markers (GSTpi) 56 days post-injury. Here, we found that this cellular response occurs earlier in rats.

Subsequently, we analyzed the origin of the OLGs that colonized the damaged area. Consistent with other studies (Zhang *et al*., 2013), our data demonstrated that oligodendrogenesis occurs in the lateral ventricle, reaching a maximum of CldU^+^/Olig2^+^ cells at 5 days post-pMCAo and later reducing at 21 days, suggesting a migration process. However, we did not find any CldU^+^/Olig2^+^ cells in either the parietal cortex or any pMCAo group. Our findings and others (Bonfanti *et al*., 2017) strongly support the notion of early recruitment of OPCs to the ischemic area through migration from surrounding regions and the differentiation of these cells into mature OLGs.

We previously described that cerebral ischemia induces an imbalance between 3RTau and 4RTau triggered by an increment in 3RTau^+^/Olig2^+^ cells in the damaged area (Villa González *et al*., 2020). It has been suggested that 3RTau promotes a more dynamic cytoskeleton (Qian and Liu, 2014). Our results demonstrated the presence of 3RTau in most pre-myelinating and some mature OLGs. This observation could be indicative of the need for OLGs to have less rigid microtubule networks, allowing them to elongate their processes and find denuded axons, as a step prior to remyelination (Gorath *et al*., 2001; LoPresti, 2002). However, it has been described that Tau is not essential for correct myelination during development as Tau-KO mice show normal myelin-levels (Torii *et al*., 2023), but rather it could be an important factor for remyelination after damage in the adult brain. Accordingly, remyelinating OLGs are not subjected to the same environment as myelinating OLGs during development (Franklin and Ffrench-Constant, 2008). This observation thus suggests that these two processes require distinct factors and are carried out differently. Tau has been proposed as a novel marker of mature OLGs (Torii *et al*., 2023). However, using *in vivo* and *in vitro* models of rat and mouse, we demonstrated the presence of 3RTau isoform in pre-myelinating OLGs, as suggested previously (Villa González *et al*., 2020). However, the existence of Tau isoforms with functional differences related to cytoskeleton dynamics and the importance of modulating the dynamic properties of OLG microtubule networks during myelination (LoPresti, 2002), prompted us to analyze the specific Tau isoforms in OLGs, with the aim to understand their potential role in remyelination after damage. Therefore, 3RTau could be a good marker of early stages of remyelination as it is expressed by pre-myelinating OLGs and mature ones.

In the period of 2 and 5 days after pMCAo the reduction in the percentage of positive Black-Gold area and the levels of MAG and MBP, well-known markers of myelin (Aboul-Enein *et al*., 2003), revealed demyelination. These results are supported by microstructural analysis with TEM that showed significant decrease in the percentage of myelinated axons and an increase in average g-ratio. This ongoing demyelination could result from the significant loss of mature OLGs in the damaged area 5 days post-pMCAo. Consistent with our results in grey matter, demyelination has been described after hypoxia in white matter (Aboul-Enein *et al*., 2003; Shi *et al*., 2015). We also observed a reduction in average axon caliber 5 days post-pMCAO, which that could be attributed to low MAG levels, a notion supported by results reported in MAG null mutation mice (Yin *et al*., 1998). Interestingly, the largest axons presented a lower g-ratio, which suggests higher vulnerability to demyelination (Liu *et al*., 2012). MOG, a minor component of CNS myelin, has been proposed to be a key player in demyelination (Dyer and Matthieu, 1994) since its location on the outermost surface of myelin makes it a target for autoimmune response, causing demyelination, and its activation has been associated with MBP degradation (Johns and Bernard, 1999). Accordingly, our results demonstrated an increase in MOG levels 5 days after damage, accompanied by severe demyelination and a concurrent reduction in MBP levels. Remarkably, our data revealed an increase in 3RTau levels 5 days post-pMCAo. These findings strongly suggest a reorganization of the microtubule network of OLGs, leading to increased dynamics, since long-term activation of MOG activity causes the depolymerization of microtubules (Johns and Bernard, 1999). These findings may indicate ongoing remyelination, in which a more dynamic OLG cytoskeleton is required to facilitate the elongation of OLG processes to contact denuded axons (Gorath *et al*., 2001), supported by our long-term results.

At 21 days post-ischemia, the damaged area showed spontaneous recovery of myelination, as reflected by the restoration of Black-Gold II staining and the MAG and MBP levels. Similar results related to the recovery of MBP mRNA levels after ischemia/reperfusion have been described but in the peri-infarct area (Gregersen *et al*., 2001). The ultrastructural analysis at this time point showed a recovery of myelinated axons and average g-ratio, thereby pointing to the remyelinating capacity of mature OLGs emerging in the damaged area. This is the first observation of spontaneous remyelination in the damaged area of cortical grey matter following pMCAo, thus highlighting the importance of long-term studies to understand the natural reparative processes of the brain (Lyden *et al*., 2021). Hence, we propose the rat pMCAo model as a valuable resource for future research in the field of remyelination.

However, the average axon caliber did not fully recover after damage. Analysis of the g-ratio versus axon diameter revealed that the largest axons showed an increase in g-ratio compared to the control group long-term. Examination of TEM images revealed a thinner myelin sheath with a similar electro-density to the control, thereby suggesting that the remyelination process did not fully restore myelin thickness, especially in large axons (Franklin and Ffrench-Constant, 2008). However, this partial restoration was sufficient to recover certain sensorimotor functions, as indicated by the results from the behavioral tests, despite the extension of the damaged area, as previously described (Hakon *et al*., 2023).

PLP, the most abundant component of myelin in the CNS, plays a key role in myelin compaction and physical stability. Interestingly, the presence of PLP has also been reported in OPCs, where it is believed to play a key role in their migration (Harlow *et al*., 2014). These findings provide a possible explanation for the stability in PLP levels observed across all experimental groups, which is consistent with the results of studies in dementia (Barker *et al*., 2013).

In postmortem brain sections from stroke patients, we observed a significant decrease in the number of Olig2^+^ and 3RTau^+^ cells in subcortical white matter compared to healthy donors. Additionally, we detected an increase in MBP coherency, indicating a reduction in myelin complexity. Specifically, vertically oriented axons remained relatively myelinated, while horizontally oriented ones were severely affected, observations similar to those described in a rodent model of transient MCAo (van Tilborg *et al*., 2017). Our results suggest that the OLG response observed in pMCAo rat model did not occur in these patients, leading to a lack of myelin repair. The absence of Olig2^+^ and 3RTau^+^ cell response in subcortical white matter, which could be necessary for proper myelin restoration, in patients who died from stroke, may have contributed to their fatal outcome.

Our findings demonstrate long-term spontaneous remyelination triggered after pMCAo that restores the functionality of some neuronal circuits. We propose that this response is a brain self-repair mechanism, although long-term analysis is needed to determine whether this restoration is maintained over time. Further characterizing the molecular mechanisms underlying this cellular response could be key to finding a novel oligodendrocyte-based therapeutical strategy to reduce the deleterious effects of ischemia and improve the quality of life of patients.

## Acknowledgments

We want to particularly acknowledge patients and Biobank HUB-ICO-IDIBELL (PT20/00171) integrated in the ISCIII Biobanks and Biomodels Platform and Xarxa Banc de Tumors de Catalunya (XBTC) for their collaboration. We are grateful to Dr Félix Hernandez and members of lab 208 at Centro de Biología Molecular “Severo Ochoa” (CBMSO) for thoughtful discussions during the preparation of this manuscript. The authors thankfully acknowledge technical assistance and guidance of Confocal and Electronic Microscopy Service at the CBMSO.

## ABBREVIATIONS

BCAS1: Brain enriched Myelin Associated Protein 1
IDI: Ischemic Damage Index
MAG: myelin-associated glycoprotein
MBD: Microtubule-Binding Domain
MBP: myelin basic protein
MOG: myelin oligodendrocyte glycoprotein
NG2: nerve/glial-antigen 2
OLGs: oligodendrocytes
OPCs: oligodendrocyte precursor cells
PDGFRα: Platelet-Derived Growth Factor Receptor Type α
PFA: paraformaldehyde
PLP: proteolipid protein
pMCAo: permanent middle cerebral artery occlusion).

---

**Supplementary Figure 1: Analysis temporal evolution of density of PDGFRα^+^ cells after cerebral ischemia.**

**(A)** Representative double immunofluorescence confocal images stained with PDGFRα (green) and Dapi (blue) of selected fields of parietal cortex of Sham and pMCAo animals. **(B)** Nissl-stained coronal brain section showing the selected fields indicated by a solid rectangle. Scale bar: 200µm **(C)** Graphical quantification of PDGFRα^+^ cells in the cerebral cortex, represented as cell density. The bars show the mean ±SEM (n=5-6). Unpaired t test; * p<0.05.

## Data availability

The data are available from the corresponding author upon reasonable request.

