## Supplementary figures and images for "The dynamics of oligodendrocyte populations following permanent ischemia promotes long-term spontaneous remyelination of damaged area"

### Supplemental Figure 1

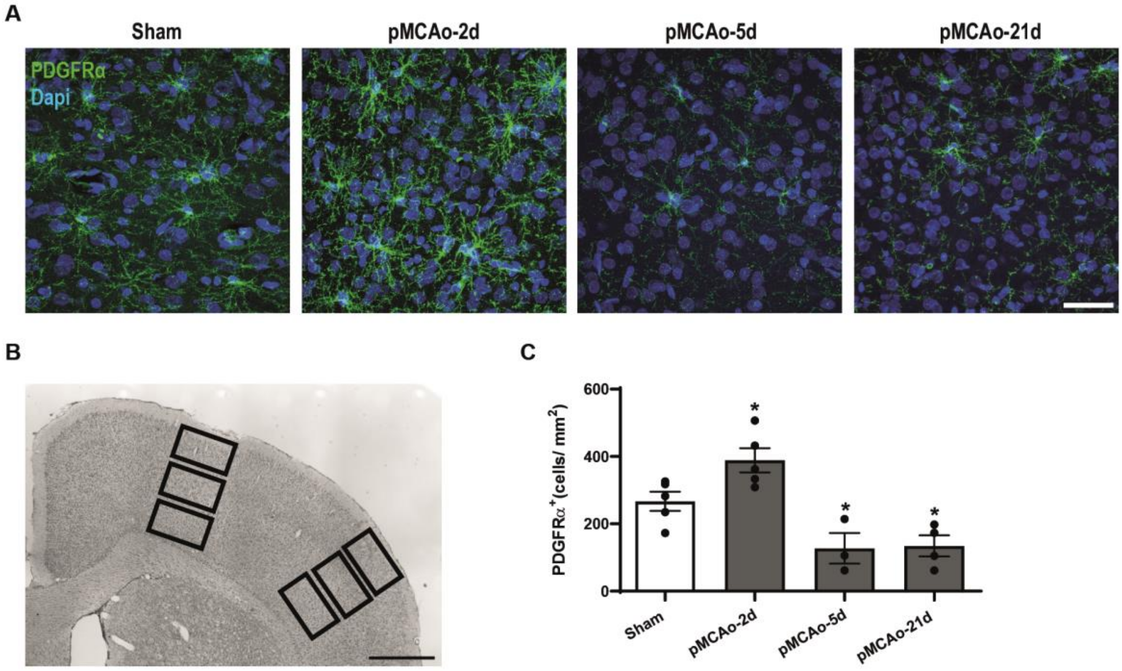


#### Supplementary Figure 1
